# False Negatives Are a Significant Feature of Next Generation Sequencing Callsets

**DOI:** 10.1101/066043

**Authors:** Dean Bobo, Mikhail Lipatov, Juan L. Rodriguez-Flores, Adam Auton, Brenna M. Henn

## Abstract

Short-read, next-generation sequencing (NGS) is now broadly used to identify rare or *de novo* mutations in population samples and disease cohorts. However, NGS data is known to be error-prone and post-processing pipelines have primarily focused on the removal of spurious mutations or “false positives” for downstream genome datasets. Less attention has been paid to characterizing the fraction of missing mutations or “false negatives” (FN). Here we interrogate several publically available human NGS autosomal variant datasets using corresponding Sanger sequencing as a truth-set. We examine both low-coverage Illumina and high-coverage Complete Genomics genomes. We show that the FN rate varies between 3%-18% and that false-positive rates are considerably lower (<3%) for publically available human genome callsets like 1000 Genomes. The FN rate is strongly dependent on calling pipeline parameters, as well as read coverage. Our results demonstrate that missing mutations are a significant feature of genomic datasets and imply additional fine-tuning of bioinformatics pipelines is needed. To address this, we design a phylogeny-aware tool [PhyloFaN] which can be used to quantify the FN rate for haploid genomic experiments, without additional generation of validation data. Using PhyloFaN on ultra-high coverage NGS data from both Illumina HiSeq and Complete Genomics platforms derived from the 1000 Genomes Project, we characterize the false negative rate in human mtDNA genomes. The false negative rate for the publically available mtDNA callsets is 17-20%, even for extremely high coverage haploid data.

## Introduction

Mutation is the process by which novel genetic variation is generated; thus, the accurate identification of mutations in genomic data is of the utmost importance for mapping Mendelian disease, population genetic analysis, tumor sequencing, and rare variant phenotype/genotype associations (Shendure and Akey 2015). Multiple bioinformatic algorithms have been developed to call mutations from short read, next-generation sequencing (NGS) data (DePristo et al. 2011; Ramu et al. 2013; Pabinger et al. 2014). However, there is a growing consensus that both short- and long-read NGS associated calling methods generate datasets with appreciably high error rates, particularly for rare or *de novo* mutations (Wall et al. 2014; Ségurel et al. 2014; O’Rawe et al. 2015). These technical error profiles affect many forms of human genomic data, and are particularly crucial for the identification of *de novo* mutations in disease phenotypes (Kong et al. 2012; Ng et al. 2010; Bamshad et al. 2011) and somatic tissue (Tomasetti et al. 2013; Costa et al. 2015). Raw 2^nd^ generation sequencing read data contains a great number of false positive variants (i.e. referred to as “sequencing error”) (Robasky et al. 2013; Reumers et al. 2011). Accordingly, pre- and post-processing pipelines filter the raw data in order to discard false positive variants. However, such pipelines may also remove true variants, which will then result in a relatively high false negative rate in the variant callset.

Recent efforts to quantify NGS error rates have primarily been focused on the identification of false positive errors in human NGS data (Zook et al. 2014; Kennedy et al. 2014). However, the need for the quantification of false negatives in such data has received far less attention (Brandt et al. 2015; Pabinger et al. 2014). High error rates complicate disease studies which search for *de novo* disease mutations between parents and probands with exome or genome sequencing. There is often a high number of candidate *de novo* mutations identified in trio/duo, but most candidates are a result of either a false positive in the offspring or a false negative in a parent (Girard et al. 2011; Veeramah et al. 2013; Vissers et al. 2010). For example, Vissers *et al*. (Vissers et al. 2010) identify 51 candidate *de novo* mutations in ten probands with mental retardation, but were only able to validate 13 with Sanger sequencing. Sanger validation of the parents revealed that only 9 of these were truly *de novo*, the remaining 4 were likely false negatives in the parents (i.e. 30% false negative rate). Other studies identify similarly high false negative rates (Michaelson et al. 2012), but the precise ratio in a given study will depend on many factors. For example, in the context of trio pedigree-based calling, filtering for mutations which are already present in a large SNP repository, such as dbSNP, will mean that recurrent *de novo* mutations are eliminated from the final callset; recent work with the EXaC database specifically highlights this problem (Lek et al. 2016). Recently, Chen *et al*. (Chen et al. 2016) report that damage introduced in-vitro during NGS library preparation results in a high number of spurious variants, and estimate that this damage causes the majority of G to T transversions in 73% of large, publically available datasets (i.e. 1000G and the Cancer Genome Atlas [TCGA]). A balanced assessment of both false positive and false negative error rates is necessary for Mendelian and complex disease identification approaches, but also crucial for evolutionary studies of mutation rates (Ségurel et al. 2014).

Previous attempts to quantify the false negative rate in NGS vary widely in approach. Perhaps the most common experiment previously conducted is the sequencing of familial trios and then Sanger validating select *de novo* mutations (see above example). Similarly, comparison between monozygotic twins or multiple NGS sequencing experiments using the fraction of mutations which fail to replicate in the same or related individuals has been reported as an estimate of the FN rate. These latter two general approaches, however, may not identify issues which systematically affect variant calling pipelines, such as filter properties. Nor is FN rates specific to human genomic sequencing (Auton et al. 2012; Nevado et al. 2014; Wang et al. 2013); work on model and non-model organisms have also produced variable FN rates. For example, short-read next generation sequencing of an inbred, non-reference mouse strain was compared to traditional BAC Sanger sequencing for 16Mb of sequence; in this inbred mouse, Keane et al. (Keane et al. 2011) estimate that the false negative rate is 6.5% and that the FN rate was twice the false positive rate. The false negative rate for short indels was notably higher (20%) (Keane et al. 2011). This study exemplifies how FN identification remained significant even for the accessible regions of a homozygous organism with a well-constructed reference genome. Marsden et al. (Marsden et al. 2016) estimate a false negative rate of 8%-10% in >15x dog genomes using both a comparison to genotyping arrays and an evolutionary calculation. Other approaches include the simulation of false negatives by introducing artificial variants into the read data directly and then estimating the fraction of artificial variants that were recovered by NGS calling pipeline. The problem with this approach is that authors may report extremely low FN rates using best-case scenarios; for example, defining “callable” sites after eliminating artificial mutations that failed coverage or other filters (Keightley et al. 2014) and as such is not a true false negative rate.

We empirically measured the false negative and false positive rates from published autosomal NGS data from the 1000 Genomes Project [1000G] and Human Genome Diversity Panel [HGDP] via Sanger-based sequencing validation (Wall et al. 2008). These results indicate that the autosomal FN rate from published datasets is highly variable and significantly greater than 1%. We then present a new phylogeny-based method to identify false negative errors in haploid non-recombining callsets (like mtDNA or Y-chromosomes) *without* generating additional validation data. The input sequences used by our method must be both homologous and non-recombining so that a single, non-ambiguous phylogeny can be constructed. Our approach can be broadly used to optimize the FN rate in haploid human next-generation sequencing experiments as set by the user. We apply our method to single nucleotide variants (SNVs) in human mitochondrial NGS data for more than 2,500 individuals. We reconsider germline mutation rate estimation in the context of false negatives by identifying *de novo* mutations from 131 mother/child duos from 1000 Genomes Phase 3 Complete Genomics data. We find that many candidate *de novo* mtDNA mutations are spurious due to a combination of false positive variants identified in the child and/or missed variants (false negatives) in the mother. Our results are in general agreement with the rates that have been calculated in previous Sanger sequencing studies (Howell et al. 2003) only if we aggressively filter the dataset.

### Estimating false negative error rates using Sanger sequencing data

We verified the ubiquity of false negatives in autosomal next-generation sequence data by a more conventional approach, comparing NGS variant calls to Sanger-based sequencing. We compare public variant call datasets from Illumina and Complete Genomics genomes to an independently published Sanger-sequencing experiment (Veeramah et al. 2012), performed on the same individuals (*see below*, all samples are cell-line derived). This Sanger data consists of short 2 kilobase intervals distributed at 40 loci throughout the autosomes. These loci were previously chosen to estimate neutral genetic diversity in human populations and hence are located at some distance from genes. These Sanger data differ from prior experiments because they were not chosen merely to validate specific NGS variants, as typically occurs for most NGS validation experiments. Hence, they represent an unbiased estimate of the false negative and false positive rates. Comparable NGS and Sanger sequence data were available for 6 Mbuti (MBI), 6 Yoruba (YRI) and 16 Luhya (LWK) (Veeramah et al. 2012).

We obtained comparable next-generation sequencing data from 28 human genomes sequenced as part of the Human Genome Diversity Project (Henn et al. 2016), Complete Genomics public dataset and the 1000 Genomes Project (Durbin et al. 2010). The forty 2 kilobase intergenic Sanger sequences for 28 individuals were independently aligned to the reference genome (GRCh37) using BLAST. We designed a Perl script that used the BLAST trace-back operation (BTOP) string to generate a VCF file. For consistency, we excluded indels from this analysis so the autosomal and mitochondrial false negative rates could be compared. We ensured that both Sanger sequences (one per chromosome) had the same start and stop position and manually trimmed problem alignments if necessary (Figure 1B). We compared the Sanger sequences to single-sample called and multi-sampled called variants, as well as imputed and unimputed versions of these callsets (Table 1). In order to generate single-sample callsets for the 1000 Genomes and HGDP data, we used BAM files released from these projects and generated VCFs using GATK’s Genotyper (-stand_emit_conf 30; -stand_call_conf 30). Emit-all files were generated so depth information would be available at all sites. Sites that fall below the stand_emit_conf or stand_call_conf are still emitted but with a LowQual flag. Complete Genomics variant data were only publically available from their proprietary single-sample calling pipeline.

**Figure 1:**
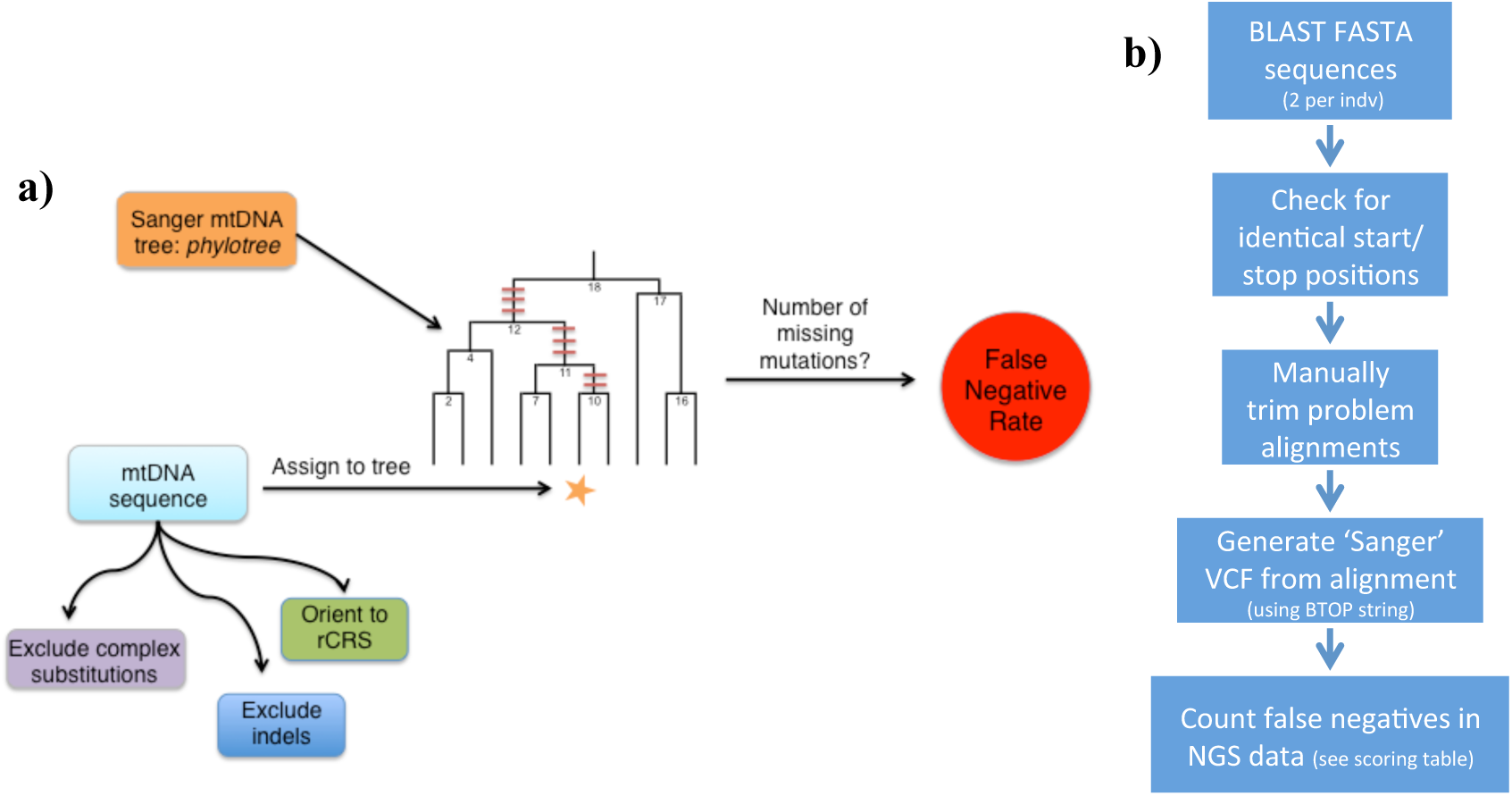
Two Schematics Illustrating False Negative Identification. A) A schematic representation of (1) the process that was used to obtain mitochondrial SNV data for each individual (before the “assign to tree” arrow) and (2) the algorithm that was used to identify false negatives in such data (*i.e.* SNVs that should have been present, but were not) based on an independently obtained phylogenetic tree (phylotree.org). B) Schematic summarizing NGS false negative identification with Sanger validation. Each box summarizes the steps and data formats that were used to identify NGS false negatives assuming that the Sanger sequenced fragments represented the true variation.

**Table 1:** *Autosomal False Negative Rates Assessed from Sanger Sequencing*

| Population | Sample ID | Multi-sample Imputed |  | Multi-sample Unimputed |  | Single-sample Calls |  |
| --- | --- | --- | --- | --- | --- | --- | --- |
|  |  | FN<sup>1</sup> | FP<sup>2</sup> | FN<sup>1</sup> | FP<sup>2</sup> | FN<sup>1</sup> | FP<sup>2</sup> |
| **LWK**<br>(1000G) | NA19027 | - | - | - | - | 12.1% | 3.30% |
|  | NA19028 | 6.0% | 0.0% | - | - | 18.8% | 2.01% |
|  | NA19041 | 8.1% | 0.7% | - | - | 14.9% | 2.70% |
|  | NA19044 | 6.3% | 1.2% | - | - | 29.9% | 2.87% |
|  | NA19046 | 5.0% | 0.6% | - | - | 32.0% | 2.87% |
|  | NA19307 | 9.3% | 0.6% | - | - | 15.4% | 1.65% |
|  | NA19308 | 7.0% | 0.0% | - | - | 4.7% | 0.58% |
|  | NA19309 | 6.0% | 0.0% | - | - | 11.1% | 1.51% |
|  | NA19317 | 4.2% | 0.6% | - | - | 18.1% | 3.01% |
|  | NA19319 | 5.1% | 0.6% | - | - | 14.2% | 1.70% |
|  | NA19346 | 3.1% | 0.0% | - | - | 19.8% | 0.62% |
|  | NA19350 | 5.3% | 0.0% | - | - | 22.3% | 0.97% |
|  | NA19360 | 3.9% | 0.6% | - | - | 12.8% | 1.67% |
|  | NA19371 | 9.6% | 2.3% | - | - | 31.5% | 3.93% |
|  | NA19373 | 6.3% | 0.0% | - | - | 21.4% | 3.14% |
|  | NA19380 | 3.9% | 1.7% | - | - | 13.7% | 3.30% |
|  | **Mean (SD)** | **5.9%** ±0.5% | **0.6%** ±0.2% | - | - | **18.3%** ±1.9% | **2.2%** ±0.3% |
| **MBI**<br>(HGDP) | HGDP00449 | 7.4% | 1.0% | 5.0% | 2.0% | 8.4% | 3.5% |
|  | HGDP00456 | 4.7% | 0.05% | 2.6% | 1.6% | 3.2% | 2.1% |
|  | HGDP00462 | 13.6% | 1.6% | 4.2% | 1.6% | 12.6% | 2.1% |
|  | HGDP00471 | 6.9% | 0.6% | 1.3% | 0.6% | 6.3% | 1.3% |
|  | HGDP00474 | 6.6% | 1.5% | 4.1% | 0.5% | 4.6% | 2.6% |
|  | HGDP00476 | 13.2% | 1.5% | 2.9% | 1.0% | 10.8% | 5.9% |
|  | **Mean (SD)** | **8.8%** ±1.5% | **1.1%** ±0.6% | **3.3%** ±0.6% | **1.2%** ±1.7% | **7.6%** ±1.5% | **2.9%** ±0.7% |
| **YRI**<br>(1000G)<sup>3</sup><br>(CG)<sup>3</sup> | NA18501 | 2.9% | 0.0% | - | - | 9.4% | 0.0% |
|  | NA18502 | 1.7% | 0.0% | - | - | 9.6% | 0.0% |
|  | NA18505 | 4.6% | 0.0% | - | - | 9.9% | 0.0% |
|  | NA18517 | 1.6% | 0.0% | - | - | 0.0% | 0.0% |
|  | NA19238 | - | - | - | - | 9.4% | 0.0% |
|  | NA19239 | - | - | - | - | 8.7% | 0.0% |
|  | **Mean (SD)** | **2.7%** ±1.4% | **0.0%** | - | - | **7.8%** ±1.6% | **0.0%** |

<sup>1</sup>Indicates the false negative rate as assessed from comparison to 80 kilobases of Sanger sequencing from the same individual.
<sup>2</sup>Indicates the false positive rate as assessed from comparison to 80 kilobases of Sanger sequencing from the same individual. We assume the Sanger sequencing does not contain spurious mutational errors.
<sup>3</sup>The multi-sample imputed column represents low-coverage data from 1000G; the single-sample column represents the high-coverage Complete Genomes data.

We count false negatives by comparing variants identified by Sanger sequencing to NGS. Each site receives a score of 0, 1, or 2: 0 indicates concordance between the Sanger and NGS genotype call, a score of 1 indicates that one alternate allele was not identified by NGS, and a score of 2 indicates that both alternate alleles were missed by NGS. To calculate the false negative rate for a sample, the scores for each site are totaled and divided by the number of expected calls, that is, the number of alternate alleles from the Sanger sequencing and multiplied by 100. Imputation is expected to identify highly probable genotypes that were absent in an initial dataset; imputation is a standard “refinement” feature of large genome datasets – including publically available 1000G and HGDP. We therefore score imputed and unimputed datasets differently, penalizing missing variants in an imputed dataset even if there is no coverage at the locus (see **Table S1**). In addition, we penalize sites in our single-sample emit-all VCFs that were called variants but marked with LowQual, as these variants would not be included in a typical variant-only file. We count false positives and the false positive rate using a similar scoring system. A score of 0 at a given site indicates concordance between the NGS and Sanger variants, a score of 1 indicates that one alternate allele was identified by NGS as variant but was reference in the Sanger data, and a score of 2 indicates that a site was called homozygous alternative by NGS and homozygous reference by Sanger.

### False Negative and False Positive Autosomal Rates for Public NGS Callsets

We first examine the single sample called variants for all three datasets. The 1000 Genomes and HGDP data represent low coverage autosomal datasets (∼7x) (*Supplementary Information, Table S1*), while Complete Genomics represents a high coverage dataset (∼54x). We expect the high coverage data to have significantly lower FN rates due to enhanced read coverage. In the 1000G LWK samples, we observe an average false negative rate of 18.3% in the unimputed single-sample callset (**Table 1**). In the HGDP Mbuti samples, the observed mean false negative rate was 7.6% in the single-sample callset. The observed Complete Genomics YRI genomes single-sample FN rate was 5.7% (**Table 1**). Our results indicate a significant difference between the single-sample calling of the 1000G false negative rate (18.3%) and the HGDP false negative rate (7.6%) which may be due to factors including library preparation, read length, sequencing instrument sensitivity, and base or variant recalibration. Both callsets were generated using parameters as specified in GATK’s best practices documentation. The coverage was marginally higher on average in the 1000G dataset (7.43x) than HGDP (6.71x). Surprisingly, the high coverage CG data had a FN rate nearly identical to the low coverage HGDP genomes. This demonstrates that *coverage alone* is not the primary determinant of FNs in a dataset; other factors, likely variant filters and sequencing technology (Robasky et al. 2013; Reumers et al. 2011) are important determinants.

False negatives are common at low coverage sites but are also observed at higher coverage sites in our dataset, e.g. 17x in a LWK individual (NA19307, chr4, position 27450119) (**Figure S2**). With biallelic sampling, the probability of sequencing only one gamete 17 times is very low (*p*=7.6*10^−6^); however, given the large number of sites obtained from whole genome sequencing the expected number of sites obtained from only a single gamete is still very large (e.g. ∼19,000 assuming even 17x coverage of 2.5Gb). Not all of these sites, however, will contain a non-reference variant. We systematically investigated the range of coverage for false negative variants identified from the single-sample called datasets (Figure 3). It is important to note that we do not call a site FN in the single-sample callset if there is 0x coverage; however, if there were sufficient reads present for the emit-all determination of a reference allele, then we do consider the site in our FN rate. We observe FNs at a range of coverages, most of the Illumina FNs were covered by 4x or more reads (Figure 3, **S2**). Coverage is far less variable in the smaller Complete Genomics dataset, but we still observe false negatives at ∼40x to almost 90x coverage. Contrary to our expectation, we find that the allele frequency for a variant is not a good proxy for estimating the probability of a false negative given there is almost an even distribution of false negatives across all allele frequencies in our sample population (**Figure S3)**.

Imputation is often used to ‘fill-in’ missing variants that may be present at appreciable frequency in a genomic dataset but are missing in a given individual due to variation in coverage or stochastic sampling. We considered the FN rates in multi-sample called and imputed datasets. In the 1000 Genomes LWK samples, we observe an average FN rate of 5.9% in the imputed multi-sample callset (Table 1). We were unable to calculate the FN in a multi-sample unimputed dataset for these LWK samples as 1000 Genomes does not make this variant dataset publically available. In the HGDP Mbuti samples that were imputed and multi-sample called, we observe an 8.8% FN rate but a 3.3% FN rate in multi-sample unimputed calls (see Table 1).

Finally, we also calculated the false positive rates for these samples using the same Sanger dataset in order to assess potential trade-offs in sensitivity versus specificity (Table 1, **Figure S4**). False positive rates were indeed much lower than FN rates, indicating that calling pipelines implemented for these public datasets were optimized for reducing false positives. The 1000G single-sample dataset has a false positive rate of 2.2% and is further reduced in the imputed multi-sample calls to 0.6%. The HGDP callsets have false positive rates of 2.9% in the single-sample callset, 1.2% in the multisample unimputed callset, and 1.1% in the multi-sample imputed dataset. We did not observe a single FP in the Complete Genomics dataset, suggesting that the CG variant calling pipeline strongly optimizes for accurate specificity of FPs.

### mtDNA FN identification

To identify false negative variants without generating additional experimental data, we leverage the phylogenetic nature of genetic sequences. In the absence of recombination, any given contiguous sequence of nucleotides can be modeled as being inherited identically by descent (IBD) by creating a phylogenetic tree of shared and derived mutations. In the absence of repeat mutation, any two DNA sequences with a recent common ancestor will share a set of mutations IBD, as well as carry their own unique and derived mutations. Using a detailed public mtDNA phylogeny (phylotree.org), we identified all variants shared by multiple mtDNA genomes such that they form the internal branches of the phylogeny. These mtDNA variants have been identified via Sanger sequencing; over the past 15 years, over 20,000 mtDNA genomes have deposited in NCBI and carefully curated by a variety of consortiums (phylotree.org, mitomap.org). We estimate the false negative rate for each sample by assigning a next-generation sequenced individual to a haplogroup in the phylogeny, and count the number of missed variants using the known set of mutations for the assigned haplogroup (see Figure 1). It is important to note that mutations on terminal branches are excluded, as we do not know whether these maybe private to the given sample used to build the tree. In addition, the rate of back mutation is assumed to be negligible but could be implemented in this model. HaploGrep was designed to be robust haplogroup assignment tool, considering the entire mitochondrial genome or any subset of it (Kloss-Brandstätter et al. 2011). As such, even if a variant that defines a large clade on the phylogenetic tree is removed, HaploGrep is still able to accurately place the genome in the proper haplogroup, albeit with a lower confidence score. Therefore, we can still use the phylogenetic nature of the mitochondrial genome to identify missing variants.

False negative estimates were calculated for two different sequencing platforms: 393 individuals in pedigree trios were sequenced by Complete Genomics (Drmanac et al. 2010) and 2,535 individuals from the 1000 Genomes Phase 3 dataset (Auton et al. 2015) sequenced via Illumina HiSeq platforms. Many individuals that were sequenced with Complete Genomics were also sequenced in the 1000 Genomes dataset allowing us to directly compare the false negative rate between the two platforms. The Complete Genomics data contained variants (up to about 50 base pairs in length) called via the company’s assembly pipeline with respect to the revised Cambridge Reference Sequence (rCRS) of the human mitochondrial genome (Andrews et al. 1999; Anderson et al. 1981). The Illumina dataset consisted of mitochondrial genomes sequenced using 75 - 100 bp paired-end reads that were mapped using the Burrows-Wheeler Aligner (BWA) software (Li and Durbin 2009). Variant calling by the 1000 Genomes Consortium was performed with the Genome Analysis Toolkit (GATK) software (DePristo et al. 2011; McKenna et al. 2010). During preparation of the callset, it was assumed that for any given locus the mtDNA has only one allele in a particular individual and heterozygous sites were removed.

We developed a software tool, PhyloFaN, that accepts VCF or VAR files as input data and is capable of lifting over variants detected using hg19 reference genome to the rCRS mitochondrial reference genome; variants detected with GRCh37 (NC_012920) do not require liftover to rCRS as the mitochondrial sequences are identical. Once input variants are aligned to match the rCRS numbering convention, both insertions and deletions are removed from the sample callset and complex multi-nucleotide polymorphisms and multi-allelic sites are split into individual records. Mitochondrial haplogroups are then assigned to each sample using the HaploGrep algorithm (Kloss-Brandstätter et al. 2011) according to the mtDNA phylogenetic tree (van Oven and Kayser 2009).

We isolate the expected internal variants for each sample based on its assigned haplogroup, and computed the rate in which haplogroup-defining variants were not observed. This rate is computed with a Bayesian inference routine, the Bernoilli *p* is used with a Jeffreys prior. The number of Bernoulli experiments, *n*, is represented as the number expected variants, given its haplogroup assignment, and the number of successes, *k*, is represented by the number of variants out of the expected that were found (see Supplemental Methods for details).

In the Complete Genomics dataset, our algorithm estimates that 2,313 out of 11,429 predicted variants were missing from the NGS variant callset. This corresponds to a false negative rate of 20.2% (confidence interval, CI: 19.5%-21.0%). We repeated the procedure for the Illumina mtDNA and obtained a false negative rate of 21.3% (95% C.I. between 21.1% and 21.5%). The ∼2,300 variants identified as missing in the Complete Genomics data and the ∼18,100 variants identified as missing in the Illumina data are plotted according to their mitochondrial base pair location (Figure 2). False negatives are particularly enriched in the hypervariable regions, despite excluding indels from the FN calculation. The tandem repeats in the hypervariable region could cause an increase in *de novo* mutations due to replication slippage, which is common in an origin of replication with a repetitive nature. However, false negatives were not enriched in regions of repetitive sequence as identified with RepeatMasker (regions shown in Figure 2). Higher mutation rate in this region could be responsible for a fraction of the apparent ‘false negatives’ observed using this model (but see below).

**Figure 2:**
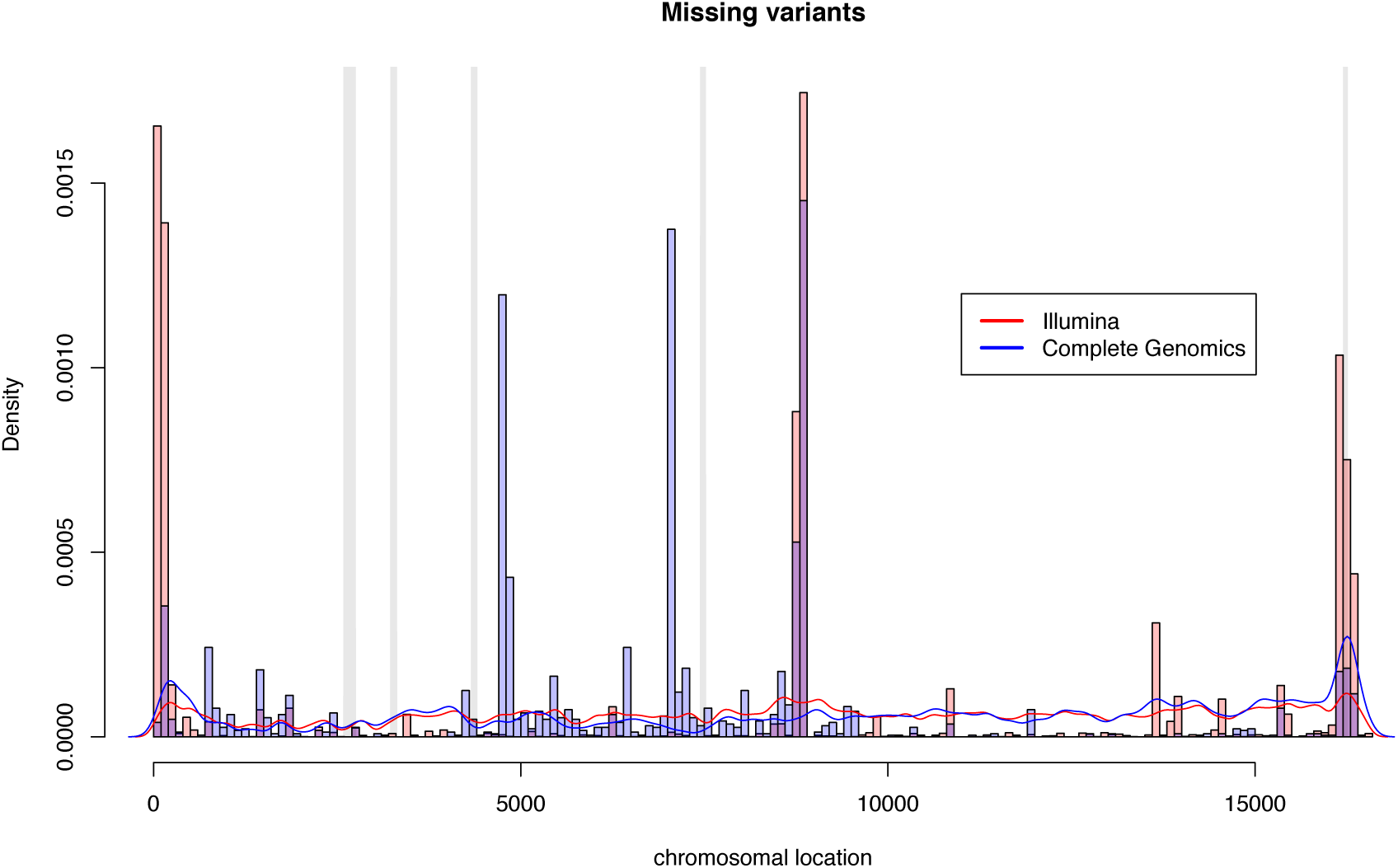
Histogram of mtDNA False Negatives by Chromosomal Location. Grey bars indicate areas of repetitive sequence on the mitochondrial chromosome, obtained by the application of RepeatMasker (including simple repeats) to the mitochondrial sequence.

**Figure 3:**
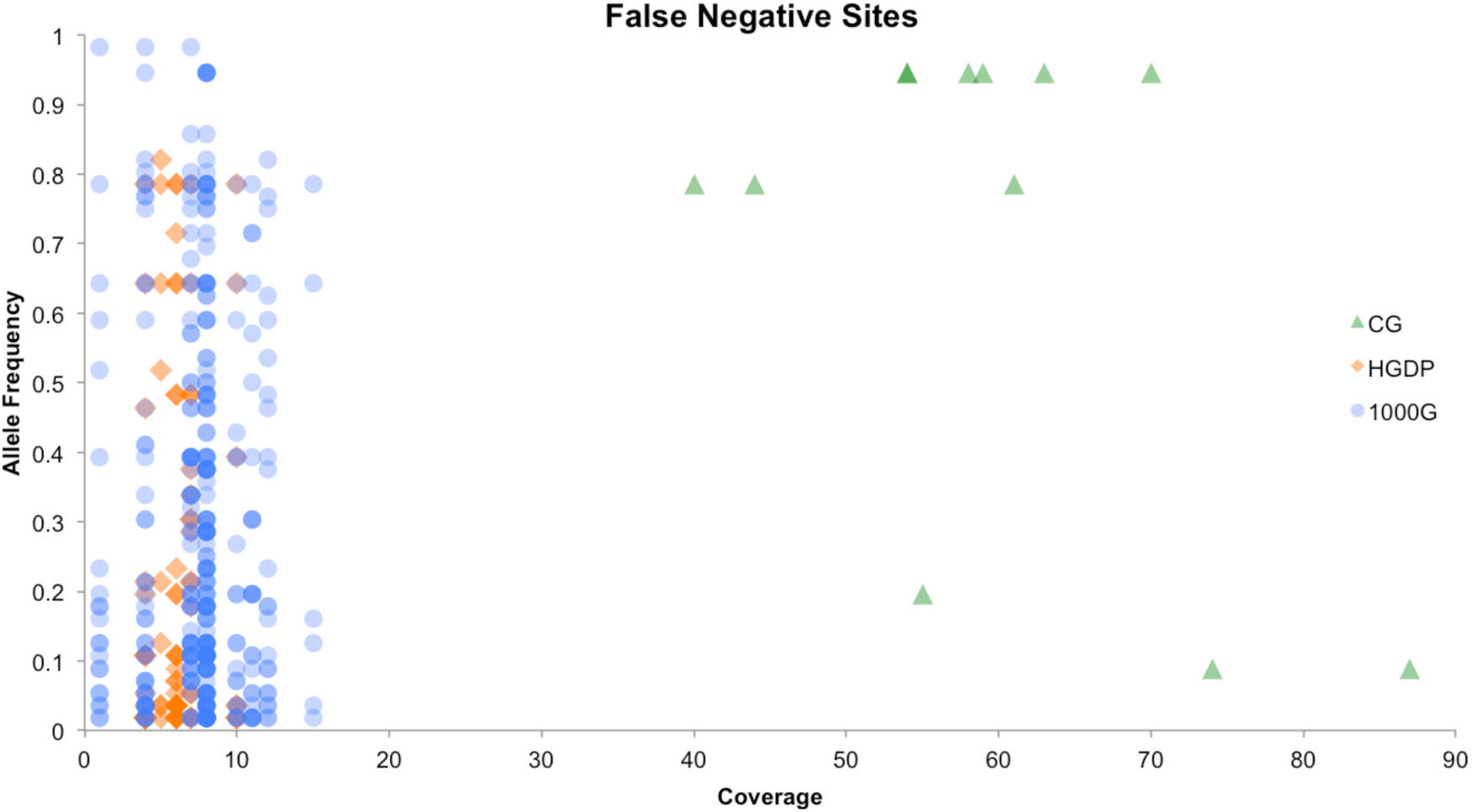
Autosomal False Negative Sites by Coverage and Allele Frequency. Each dot represents an NGS autosomal false negative (FN) site in one individual from the single-sample variant calling dataset. FNs are colored by project (Table 1). The Complete Genomics dataset has higher mean coverage than the two Illumina datasets in our study. To calculate allele frequency, Sanger data from the three African populations were combined and the allele frequency across the dataset was estimated as the non-reference allele frequency. The y-axis represents the non-reference allele frequency relative to hg19. False negatives span the full frequency range in all three datasets.

We investigated whether the loss of these putative true variants from the callset was due to the strictness of filters applied by the post-processing pipelines. Such pipelines tend to optimize filtering out false positive variants, which are highly prevalent in raw 2^nd^ generation sequencing data (DePristo et al. 2011; O’Rawe et al. 2013). We hypothesized that these pipelines would often miss large numbers of variants that are, in truth, present within the raw sequence data. We confirmed that the missing variants were indeed present in the pre-pipeline BAM files. For example, the HaploGrep algorithm assigns individual HG00097 to haplogroup T2f1a1. The mitochondrial phylogeny has a substitution from G to A at position 8860 (rs2001031) derived in the reference sequence haplogroup, H2a2. Thus, the phylogeny predicts that the mitochondrial sequence of HG00097 would to have a G at this locus, whereas the reference sequence has an A. This G variant was not present in the sample VCF file but was indeed present in the original BAM file for this sample (**Figure S1**).

The variant at location 8860 contributes to the false negative rate in our case study to a great extent due to the fact that it defines the H2a2 haplogroup of the Cambridge Reference Sequence. It is predicted but missing in 2,533 of the 2,535 Illumina VCF files. As a result, the corresponding 2,533 false negative calls comprise 17% of the false negatives we detect in the Illumina callset. This variant represents the central peak in Figure 2. We also checked whether the exclusion of multi-allelic variants in the public 1000G Illumina mtDNA files significantly affects the FN rate. Here, “multi-allelic” is defined as a locus that has more than two alleles present among the sampled individuals. As per our expectation, allowing for multi-allelic variants in the VCF improved the false negative rate, decreasing from 21.3% to 17.1% (C.I.: 16.8%-17.3%). Multi-allelic variants may be more common in mtDNA datasets than autosomal due to high relative mutation rate of mtDNA.

We further investigated the effect of decreasing the number of stringency filters in the GATK pipeline on the false negative rate. Specifically, we recalled variants using the original BAM files using GATK’s UnifiedGenotyper (McKenna et al. 2010) with respect to the GRCh37 reference sequence. The choice of reference genome is significant because most variant calling pipelines align to a reference genome from which non-reference variants are identified; FN’s are, by default, assumed to carry the reference allele (Degner et al. 2009; Brandt et al. 2015). We ran GATK default emit-all parameters (i.e. permissive genotype calling) on each individual sample separately (referred to as “single-sample calling”). This pipeline dramatically reduced the false negative rate in Illumina data, from 17.1% down to 2.28%. We note that is approach is likely associated with a great number of false positive variant calls because these filters are built to minimize sequencing error in autosomal sequences – but the experiment demonstrates how the majority of the false negatives are present in the short read data and erroneously excluded in subsequent filtering processes. In summary, we find that most FN variants are missing due to filtering within the post-processing pipeline.

We then tested whether false negative status of a variant correlates with short read coverage. To compute the depth of coverage for each base pair location in each sample in our Illumina data, we used GATK’s DepthOfCoverage (McKenna et al. 2010). Average coverage for the variants identified as false negatives was found to be 2,022x, while that for the variants expected from the phylogeny and contained in the VCF files was 2,244x. The difference between coverage values of the former group and those of the latter group was shown to be significant by a t-test (p < 2.2×10^−16^, 95% C.I. for the difference in coverage between 197x and 248x). A best-fit logit linear regression model of the dependence of false negative status on coverage based on the same data had an intercept equal to −1.36 and a coefficient of the explanatory variable equal to −0.000105. A logit model with these parameters predicts that an increase in coverage from 2,000 to 3,000 reads leads to a decrease in the probability of false negative status from 17.3% to 15.8%. In other words, for our extremely high-coverage data, the effect of coverage on the false negative error status via the pipeline filters is significant but not very large.

### mtDNA de novo mutation rate

Our results indicate that the false negative rate is sensitive to variant filtering and coverage, leading to false negative rates of 6-20% for publically available datasets. We illustrate the effect of the FN rate by considering its impact on estimates of the human mtDNA germline mutation rate. We estimate the germline mtDNA mutation rate by comparing mitochondrial genomes of mothers with those of their children to identify *de novo* mutations. We utilize 131 mother-child pairs from the 1000 Genomes Phase 3 Complete Genomics samples (Auton et al. 2015) for which genotype calls were provided by proprietary CG processing pipelines. We identified 36 mitochondrial SNVs that were present in the child but not in the mother (Table 2). For each mother-child pair that had a putative *de novo* mutation, we used HaploGrep to assign haplogroups on the basis of the known mitochondrial phylogenetic tree (Kloss-Brandstätter et al. 2011; van Oven and Kayser 2009), separately for both the mother and the child.

**Table 2:** *Candidate de novo mitochondrial mutations from 131 duos*

| Mother ID | Child ID | Source | Haplogroup Assignment | Candidate *de novo* mutation<sup>1</sup> | FP in child due to 'No call' in Mother<sup>2</sup> | FP in child due to FN in Mother<sup>3</sup> |
| --- | --- | --- | --- | --- | --- | --- |
| GS000016020 | GS000016048 | LCL | U4b1b1 | 8655 (242) |  |  |
|  |  |  |  | 10566 (22) |  |  |
| GS000016039 | GS000016539 | LCL | H2a5b / H2a5 | **9835** (8342) | 3166 (7413) |  |
| GS000016041 | GS000016538 | LCL | U5b2a1a+16311 / U5b2 | **204** (9262) |  |  |
| GS000016398 | GS000016412 | LCL | T2 / T2b | 14050 (629) |  |  |
| GS000016456 | GS000016408 | LCL | H52 | 2351 (2147) |  |  |
| GS000016465 | GS000016380 | LCL | H5b1 | **279** (13042) |  |  |
|  |  |  |  | **14569** (8028) |  |  |
| GS000017172 | GS000017223 | LCL | L3e2b1a2 | 2045 (1342) | 2483 (73) | 16189 (516, L3e2b) |
| GS000017130 | GS000017271 | Blood | B2 | 3173 (2494) |  |  |
| GS000016414 | GS000016400 | LCL | U5a1a2b |  | 750 (2424) | 1700 (6051, U5a1a) |
|  |  |  |  |  | 1438 (1584) | 3197 (17404, U5a'b) |
|  |  |  |  |  | 2706 (10860) | 11467 (6271, U) |
|  |  |  |  |  | 10915 (8882) | 14793 (8883, U5a) |
|  |  |  |  |  | 14766 (4244) | 15218 (8736, U5a1) |
|  |  |  |  |  | 15326 (10230) |  |
| GS000016469 | GS000016540 | LCL | H3a1a |  | 4769 (129) |  |
| GS000017276 | GS000017275 | Buffy | L1b1a15 |  | 3936 (1003) |  |
| GS000016396 | GS000016409 | LCL | U3a1a/U3a1 |  | 8860 (205) |  |
| GS000016011 | GS000016459 | LCL | W1c1 |  | 2706 (12255) |  |
|  |  |  |  |  | 4769 (305) |  |
| GS000017242 | GS000016026 | LCL | U5a1 |  | 2706 (11788) |  |
|  |  |  |  |  | 8994 (1864) |  |
| GS000017185 | GS000017173 | LCL | L2a1f |  |  | 16192 (4551, L2a1f) |
| GS000017227 | GS000017119 | LCL | L2b3a |  |  | 16213 (6945, L2b) |
| GS000017045 | GS000017047 | Buffy | L1b1a15 |  |  | 16355 (6228, L1b1a15) |
|  |  |  | **Total count** | **10** | **15** | **9** |

<sup>1</sup> Mitochondrial base pair position for each candidate *de novo* mutation which appeared in the child but not the mother. The variant quality score for the child's SNV is indicated in parentheses. Quality scores greater than 5000 are indicated in bold (see supplemental Figure S3 for bimodal distribution of variant quality scores).
<sup>2</sup> mtDNA position of candidate *de novo* mutations which were inferred to be false positives due to a 'no call' in the mother. The variant quality score for the child's SNV is indicated in parentheses.
<sup>3</sup> mtDNA position of candidate *de novo* mutations which were inferred to be false positives in the child inferred from the local mtDNA phylogeny; mother's allele was indicated as reference but the mutation was derived in the derived haplogroup (i.e. a haplogroup defining mutation). Haplogroup defined in parentheses.

Next, to confirm maternal assignment for each duo, we compared the haplogroups that HaploGrep assigned to the mother and the child for each of the pairs. We retain pairs for which either (a) the mother’s haplogroup was the same as the child’s, (b) the child’s haplogroup was contained within the mother’s or (c) the mother’s haplogroup was contained within the child’s. Two of the 34 candidate *de novo* mutations came from mother-child pairs whose haplogroup pairs did not fall in either of the above-mentioned categories indicating sample swaps or non-maternity. For each of the remaining candidate mutations, we then checked the variant call in the mother’s data. We excluded 15 candidate *de novo* mutations that had been identified within segments that had a ‘no-call’ status in the mother (Table 2). We then applied our phylogenetic method of identifying false negatives to each of the remaining 19 candidate *de novo* mutations to test whether the absence of each of these variants in the mother’s sequence was due to a false negative call at the locus in question. For 9 of the remaining candidate mutations, the variants in the mother’s sequence were predicted to be present based on the mother’s phylogenetic lineage, so the corresponding candidate mutations were excluded. This left us with 10 candidate *de novo* mutations: 8 in the coding region and 2 in the control region.

In order to assess whether our final *de novo* mutation candidates could still be false positives, we compared their variant quality scores (varScoreVAF column in the Complete Genomics VAR files for children that had the corresponding SNVs) with those of all SNVs identified in our Complete Genomics data (**Figure S5**). The ten *de novo* mutation candidates have significantly lower variant quality scores than the rest of the SNVs in the dataset (Mann-Whitney test, *p* = 0.02) indicating that many of them are likely spurious variants. Only 4 variants had a varScoreVAF (derived from maximum likelihood model) greater than 5000.

Using the 131 duo dataset, we could then compare the NGS mtDNA coding region mutation rate with earlier Sanger sequencing estimates. By dividing the number of putative *de novo* mutations by the total called sequence length (see *Supplemental Methods*), we obtain an estimate of 8 / 1,981,090 = 4.04×10^−6^ mutations per base pair per generation (95% CI: 1.74×10^−6^ − 7.96×10^−6^) in the human mitochondrial coding region and 2 / 146,610 = 1.36×10^−5^ mutations per base pair per generation in the control region (95% CI: 1.65×10^−6^ and 4.93×10^−5^). However, if we consider only the two coding region mutations which have a quality score greater than 5000, then we obtain an estimate of μ = 1.01×10^−6^/bp/g. Howell and colleagues (Howell et al. 2003) produce a Sanger pedigree-based estimate of the coding region mutation rate of 6.0×10^−7^ bp/g (99.5% CI: 8.0×10^−8^ −2.0×10^−6^ bp/g), after conversion from the divergence rate to the mutation rate and assuming 20 years per generation; inclusion of all 8 mutations in Table 1 results in a mutation rate outside of the earlier confidence interval, while our more conservative 1.01×10^−6^/bp/g is consistent with Howell et al.’s meta-analysis. A recent mtDNA genome μ estimate of 2.7×10^−7^/bp/g from extremely high coverage NGS sequencing of 39 mother-child duos is significantly lower than the pedigree estimate here; however Rebolledo-Jaramillo et al. (Rebolledo-Jaramillo et al. 2014) note that they heavily filtered variant calls in order to confidently discriminate heteroplasmies from sequencing artifacts, and therefore their estimate should be seen as a lower bound. We note that multi-generational pedigrees are needed to discriminate *de novo* germline mutations from somatic mutations. Assuming that the true FN detection rate in the CG dataset is ∼20%, one could also argue that our μ should be corrected by 20% to account for missing *de novo* variant calls in the child (**Table S2**).

### Conclusions

We demonstrate that false negative mutations are a significant feature of short-read, next-generation sequencing data sets. While various previous reports have estimated FN rates for specific datasets, the general assumption is either that read coverage is the primary determinant of false negatives or imputation will correct for sparsely missing variants. Gross characterization of the expected false negative rate for any given sequencing experiment is often difficult because the FN rate is sensitive to post-processing variant calling pipeline parameters. Previously, these pipeline parameters have been optimized for false positive removal (Table 1). Autosomal FN results presented here range between 3%-18% for large, publically-available human genome datasets. While 80 total kilobases per individual contains relatively few SNPs, our validation approach here is unique for using an unbiased dataset that was not chosen to specifically validate *de novos* for independent published datasets.

We provide a computational tool [PhyloFaN] for rapid estimation of the FN rate in new genomic datasets, which will allow optimization of the FN rate without relying on new Sanger sequencing validation. Our phylogenetic approach is currently confined to non-recombining homologous loci, such as mtDNA and the Y chromosome. While PhyloFaN can be used to systematically explore the effect of pipeline parameters on the false negative in haploid systems, it is an imperfect proxy for assaying autosomal data. Future extensions using ancestral recombination graphs or ARGs (Rasmussen et al. 2014), however, hold great potential. Finally, we describe one instance in which increasing coverage hundredfold still results in large false negative rates by exploring FNs and FPs in 131 high-coverage mtDNA duos from 1000 Genomes. We find that the majority (89%) of putative *de novos* identified in the child are due to either variant quality issues or false negatives in the mother (resulting in a ‘false positive’ *de novo* for the child). Accurate identification of *de novo* mutations remains a critical challenge for Mendelian disease, cancer genomics and mutation rate estimation.

## Acknowledgements

We thank Krishna Veeramah for providing the Sanger sequencing dataset. We thank Adam Siepel, Jeffrey Kidd and Vagheesh Narasimhan for valuable discussion. Finally, we are grateful to Jason O’Rawe for coding the mtDNA false negative pipeline. D.B. was supported by The Research Foundation at SUNY Stony Brook.

